# Integrated analysis of Wnt signalling system component gene expression

**DOI:** 10.1101/2022.03.07.483288

**Authors:** Paula Murphy, Chris Armit, Bill Hill, S. Venkataraman, Patrick Frankel, Richard Baldock, Duncan Davidson

## Abstract

Wnt signalling controls patterning and differentiation across many tissues and organs of the developing embryo via temporally and spatially restricted expression of multi-gene families encoding ligands, receptors, pathway modulators and intracellular components. Here we report an integrated analysis of key encoding genes in the 3D space of the mouse embryo across multiple stages of development. We applied a method for 3D/3D image transformation to map all gene expression patterns to a single reference embryo for each stage providing both visual analysis and volumetric mapping allowing computational methods to interrogate the combined expression patterns. We identify novel territories where multiple Wnt and Fzd genes are co-expressed and cross-compare all patterns, including all seven Wnt paralogous gene pairs. The comprehensive analysis allows regions in the embryo where no Wnt or Fzd gene expression is detected, and where single Wnt genes are uniquely expressed, to be revealed. This work provides insight into a level of organisation of the patterns not previously possible, as well as presenting a resource that can be utilised further by the research community for whole system analysis.

## INTRODUCTION

The Wnt signalling system of cell-cell communication is ancient and fundamental to the construction of an organised animal body plan (Loh et al., 2016), proposed to have arisen concurrently with the metazoan lineage (Moroz et al., 2014; Ryan et al., 2013). Since the original dual discoveries of key roles for Wnt in development and dysregulation during oncogenic transformation, it has more recently been shown to control cell differentiation within, and maintenance of, stem cell niches (Clevers et al., 2014). Spatio-temporally localised Wnt signalling plays a key role in patterning the primary body axis across very different body plans (Holstein, 2012) and is required for the establishment and healthy maintenance of organ systems from the Central Nervous System (CNS) to the kidney and gut (Noelanders and Vleminckx, 2017; Tian et al., 2018; Wang et al., 2018). Clearly, the spatio-temporal expression of genes that control Wnt signalling is of central importance.

The Wnt system is complex, with components encoded by several highly conserved multi-gene families; for example, there are 19 conserved Wnt ligand encoding genes in all mammals, each uniquely essential as demonstrated by mutation analysis in the mouse and the level of sequence conservation between species. 10 genes encode Frizzled (Fzd) receptors, that work together with a variety of co-receptors such as Lrps, Ryk and Ror. Extra-cellular modulators of the pathways include Secreted Frizzled Related Proteins (Sfrps), Wif and Wise. Intracellularly Wnts can trigger a number of different pathways, the best understood of which is the canonical/β-catenin dependent pathway where stabilisation of β-catenin leads to gene expression changes in the responding cell; pivotal components including the Tcf/Lef transcription factors and β-catenin itself. Many other proteins interact at multiple levels in alternative pathways, influencing cellular outputs and this is rendered yet more complex through cross talk between the pathways and other key signals such as BMP (Singh et al., 2018).

A central question is, why are there so many Wnt and Fzd genes in a single organism? Indeed, there are 12 sub-families of Wnt genes conserved across metazoans, with gene loss and duplication in particular lineages (Somorjai et al., 2018). One possibility is that the activities of different members have become segregated during evolution to function in different spatio-temporal contexts in the developing embryo. Here we address this question by making detailed comparisons of comprehensive gene expression patterns using the approach provided by the Mouse Atlas Project which is based on the idea of mapping gene expression and other data onto a series of digital reference models of the mouse at successive stages of development (Davidson and Baldock, 2001). These reference models provide a framework in which the data can be interrogated computationally, integrated in a database with other gene expression data and, importantly, visualised in numerous ways to examine 3D spatial relations in an anatomical context (Armit et al., 2017).

We previously reported comprehensive 3D expression of Wnt and Fzd encoding genes at E11.5 (Theiler stage 19 (Theiler, 1989) (Summerhurst et al., 2008) and of Tcf/Lef transcription factors across time (Vendrell et al., 2009). Here we extend this work using the Mouse Atlas approach to examine and compare RNA expression patterns of genes encoding Wnt ligands (19 genes), Fzd receptors (10 genes), Tcf/Lef transcription factors (4 genes), Secreted Frizzled Related Proteins (Sfrp) (5 genes) and other modulatory proteins, Wif1 and Wise, as well as canonical pathway activity revealed through a reporter mouse line TCF/Lef:H2B-GFP (Ferrer-Vaquer et al., 2010). We studied three key stages when the body plan is being elaborated and various organ systems established: Theiler Stages (TS) 15 (Embryonic day (E) 9.5), 17 (E10.5) and 19 (E11.5). This approach can be used to pose questions not possible in any other way such as asking where no Wnt expression is detected or where specific groups of components are co-expressed or relate to territories of canonical pathway read-out. We set out to test the hypothesis that Wnt and Fzd expression is a mosaic of domains in which only one or a few Wnts and Fzds are expressed in each. Our results also provide insight into the similarity and divergence of expression of different Wnt genes in the mouse, including the deployment of the more recently duplicated paralogous genes and the relation between Wnt pathway component gene expression and canonical pathway activity.

## RESULTS

### Integrated visualisation of Wnt pathway component gene expression patterns across E9.5 to E11.5 in the mouse embryo

3D expression patterns for all Wnt, Fzd, Tcf/Lef, Sfrp, Wif1 and Wise genes as well as canonical Wnt pathway readout (Ferrer-Vaquer et al., 2010) were mapped onto reference embryos at E9.5, E10.5 and E11.5 (both data and reference embryos were staged precisely to Theiler Stages 15, 17 and 19 respectively (Theiler, 1989)). All data are available through University of Edinburgh *DataShare* (Murphy et al., 2021a; Murphy et al., 2021b)(https://doi.org/10.7488/ds/3141; https://doi.org/10.7488/ds/3142) and can also be viewed using an on-line 3D section viewer such as the Mouse Atlas IIP viewer (Armit et al 2015; Husz et al 2012 available at www.emouseatlas.org/WntAnalysis) or downloaded and viewed using ITKSnap (Yushkevich et al., 2006). Examples of mapped and integrated data are shown for Wnt genes (Fig. 1; movies S1-5). The accuracy of mapping (Fig. S1) has enabled an integrated analysis to reveal higher order patterns.

**Fig. 1:**
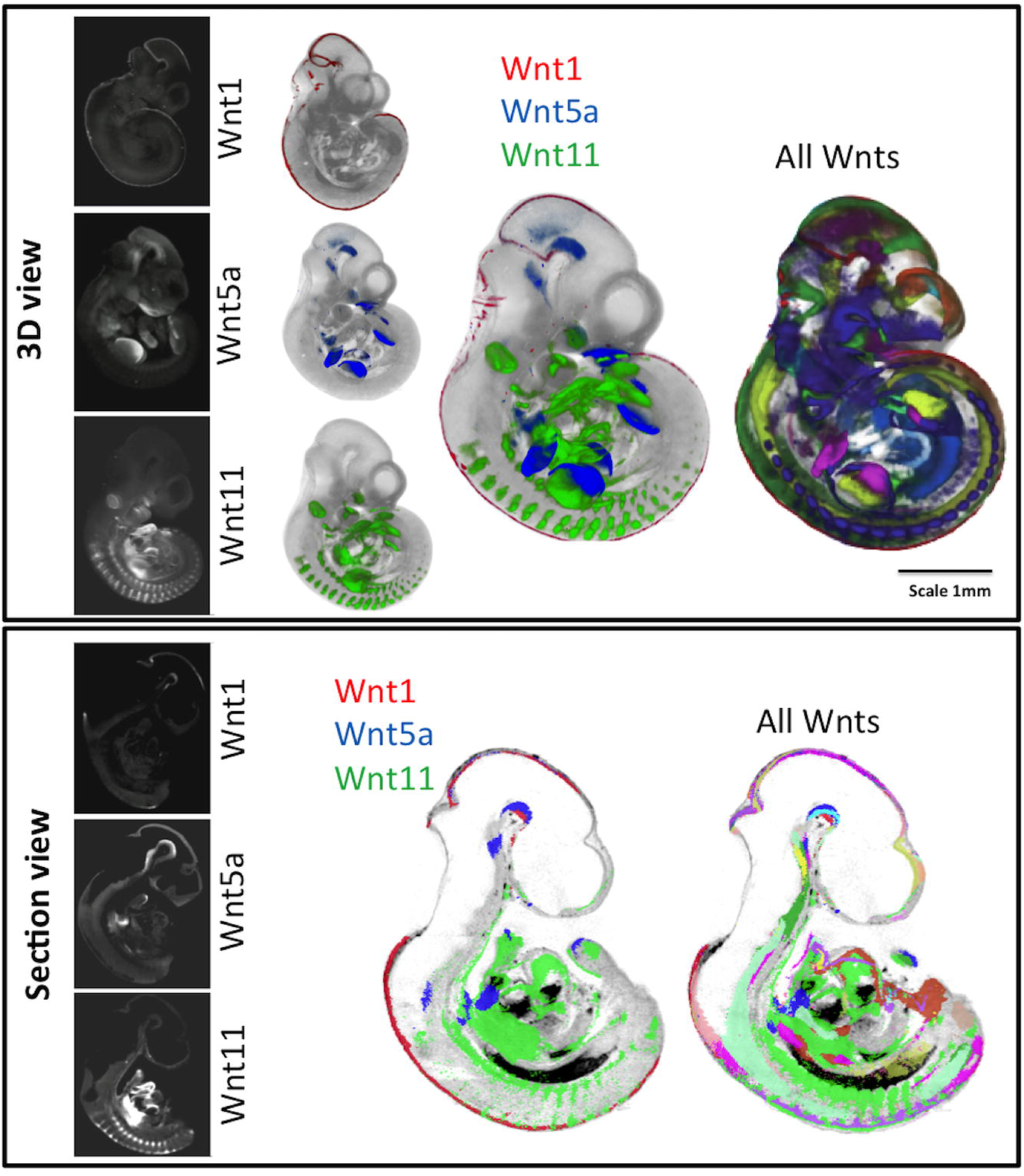
Visualisation of mapped and integrated gene expression patterns, exemplified at E10.5. Upper panel; whole embryo external views of 3D data. The original OPT reconstructions showing the expression of Wnt1, Wnt5a and Wnt11 are on the left and individually mapped to the same reference model in the next column (see full 3D movies of each in Supplementary data movies S1-3). The right-hand columns show the three patterns integrated and all Wnt expression patterns integrated (see 3D movies S4, S5). The lower panel shows virtual sagittal section views through the same 3D data, showing original OPT data and mapped data as above. Red; Wnt1, green; Wnt11, blue; Wnt5a in the middle columns. For more detail on visualization of each pattern see Movie S5.

Virtual sections through the original, unmapped OPT data have been examined on a gene-by-gene basis to complement this analysis, for example to distinguish epithelial-specific expression. Supplementary movies S1-3 show mapped domains of individual gene expression patterns at E10.5, shown in Fig. 1, while movies S4 and S5 respectively show the integration of three Wnt genes and of all Wnt genes added incrementally.

Fig. 2A shows combined domains for all Wnt ligand genes (red(i)), Fzd receptor genes (green(ii)), and canonical pathway readout from transgenic line TCF/Lef:H2B-GFP (referred to as Tcf/Lef-GFP throughout) (purple (iii)) across stages. Overlaying combined Fzd on Wnt patterns (iv) shows the overlap (yellow). Across stages, combined Fzd domains are more restricted than combined Wnt domains, so that some regions express Wnt genes in the absence of detected Fzd expression (red(iv)). These are extensive, largely ventral and visceral and notably do not overlap with canonical pathway read-out (v). In contrast, domains that show detection of Fzd expression without Wnt expression (green(iv)) are more restricted but with some notable examples such as the anterior telencephalon at E10.5, where Fzd3, 10, 6, 7 and 8 are expressed, the ventral diencephalon at E10.5 and 11.5, and part of the developing limbs at E10.5, contributed to largely by Fzd10. As expected, canonical readout, at any stage, largely fits within the overlap of Fzd and Wnt domains (v), in the dorsal aspects of the main body axis, prominent in the neural tube, and also in the limb, branchial arches and heart. We noted surprising instances where Tcf/Lef-GFP is detected in the absence of concurrently detected Wnt or Fzd expression, notably the nasal epithelium at E11.5, proximal mesenchyme of the 1^st^ branchial arch at E10.5 and E11.5 (Fig. S2) and regions in the ventral diencephalon at E10.5 and E11.5.

**Fig. 2.**
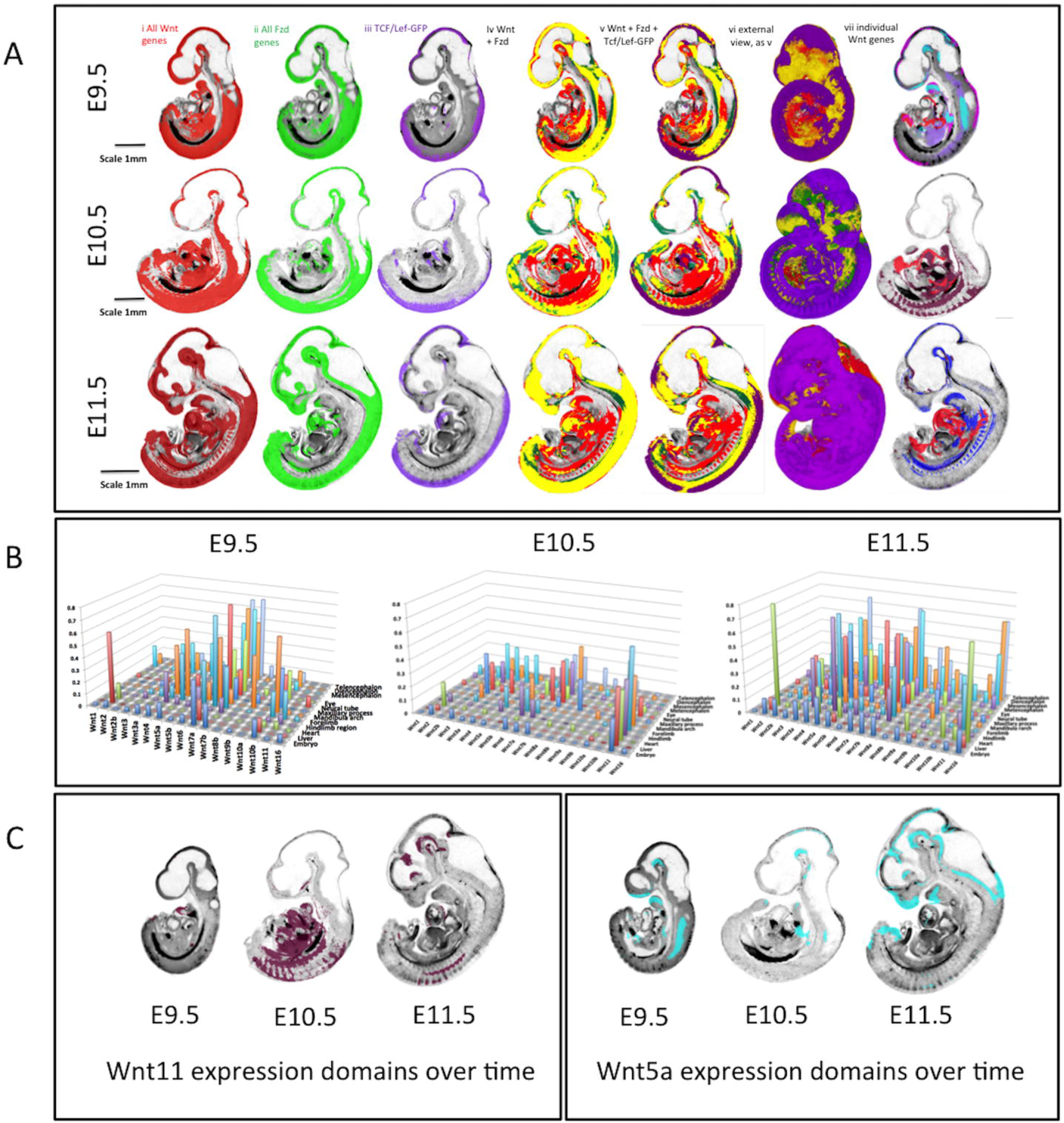
Overview of integrated expression patterns. 2A shows union of the expression of all Wnt genes (red i) all Fzd genes (green ii) and the canonical pathway readout reporter (purple iii) across the three stages of development as indicated. Column iv shows the all Fzd domain overlaid on the all Wnt domain with overlap shown in yellow. Columns v and vi add the canonical read-out domain in purple (vi is an external view). Column vii shows individual Wnt expression patterns that contribute to the ventral Wnt domain (red; Wnt2, purple; Wnt10b, pink; Wnt4, pale blue; Wnt5a, dark red; Wnt11, dark blue; Wnt5b). 2B presents 3D graphs showing the extent (proportional size) of each Wnt gene domain in the whole embryo and in individual anatomical domains across stages as indicated; the y-axis shows the proportion of the anatomical domain (z axis) occupied by each gene expression domain (x axis). 2C shows Wnt11 and Wnt5a expression domains on midline sagittal sections across stages; these patterns illustrate the dynamic changes in extent of expression across stages

Since all expression domains for each gene at each stage are digitally mapped, it is possible to analyse the domains in the context of the whole embryo or within anatomical subdomains (Baldock et al., 2003; Brune et al., 1999) (Fig. 2B, Table S1). Therefore, one can computationally ask the proportion of the embryo, or of a delineated anatomical structure such as the neural tube, occupied by an expression domain over time. For example, at E10.5, the most broadly expressed Wnt gene in the embryo is Wnt11 (22%), whereas Wnt1, Wnt8a and 8b are very restricted (Fig. 2B; ≤1%). The extent of expression of Wnt genes is dynamic. Overall expression domains are more restricted at E10.5 compared to E9.5, becoming more expansive again at E11.5 (Fig. 2B), Wnt5a being a striking example (Fig. 2C), but there are notable exceptions, including Wnt11 Wnt6, Wnt3, Wnt2 and Wnt10a, that expand expression domains between E9.5 and E10.5, to become more restricted again at E11.5 (Fig. 2C).

Comprehensive mapping shows territories where no Wnt and Fzd expression is detected (Fig. 3A; see www.emouseatlas.org/WntAnalysis). These territories are largely ventral and visceral and are broadly similar across stages. Territories where expression of a *single* Wnt gene is uniquely detected (Fig. 3B) are mostly represented in ventral, visceral regions, but also in parts of the nervous system; neural tube and diencephalon. These territories are similar between E9.5 and E10.5 but become noticeably more restricted at E11.5 (Fig. 3B). To ask which Wnts are expressed in these single-gene domains, we used parallel co-ordinate visualisation (Moustafa, 2011) across time, showing that a major contributor is Wnt2 at all stages (8, 11 and 17% across stages; representing 49, 54 and 37% of the Wnt2 domains; Table S2). Other major contributors are more dynamic (Fig. 3C). At E9.5 and E10.5, approximately one third of canonical Wnt readout domains are contained within unique Wnt expression territories, becoming about one fifth at E11.5.

**Fig. 3:**
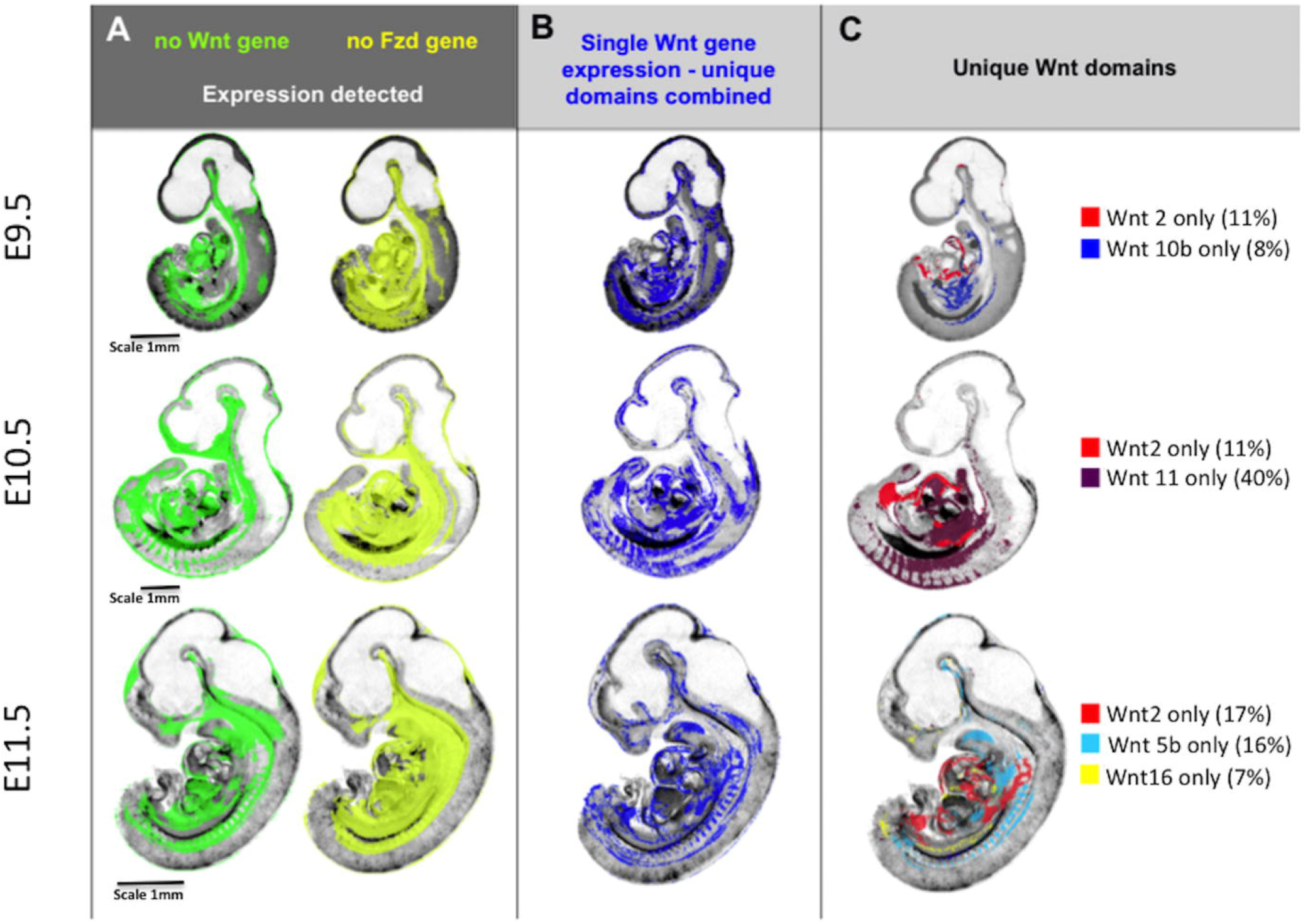
Integrative mapping of all Wnt and Fzd domains allows visualization of the territories where no Wnt or Fzd gene is expressed (A) or where unique Wnt genes are expressed (B and C). A) domains across stages where no Wnt expression (green) or no Fzd expression (yellow) is detected. B) domains across stages where a single Wnt gene is detected i.e. unique detection of a single Wnt gene transcript. C) individual Wnt gene domains that account for much of the single Wnt gene domain at each stage i.e. much of the unique Wnt gene territory in the ventral embryo at E10.5 is occupied by Wnt2 and Wnt11 expression domains whereas Wnt11 contributes little at E11.5 when Wnt5b is more prominent. The figures noted in brackets are the percentage of the unique Wnt gene expression domain at that stage contributed to by that gene. Note that the unique Wnt domains reported here were obtained by subtraction of multiple mapped expression domains and, as such, are sensitive to cumulate effects of noise in the data for each gene, in particular small differences in thresholding the original, continuously variable signals into binary (expressed versus not detected) values. While the images show the general location of the domains, the boundaries should be considered as approximate.

### Co-expression of multiple Wnt or Fzd genes

At the stages examined, most of the embryo displays the expression of 0 - 2 Wnt or 0 - 2 Fzd genes. However, certain localised regions express a notably large number of Wnt or Fzd genes where the expression of 4 - 11 Wnts or 4 - 8 Fzds maps to each image voxel in the reference model. We refer to the number of genes with expression mapped to a voxel as the *occupancy* of that voxel.

Generally, regions with high occupancy of Wnt or Fzd expression are distinct from one another (Fig. 4), each with one or a few peaks of occupancy (e.g. Fig. 4B E11.5). At all three stages, Wnt occupancy levels 1 and 2 are widely dispersed, but most level 3 Wnt occupancy domains and almost every domain of occupancy 4 and above, contains, or abuts, a region with occupancy 5 or greater (Fig. 4B). In several locations, gradients of Wnt occupancy 4 and above are steep. We have distinguished ‘regions of high occupancy’ (ROHOs) computationally as territories with occupancy above a certain threshold. For Wnt genes we used five or more (5+) at E9.5 and E11.5 and 4+ at E10.5. For Fzd genes we used 4+ at all three stages. ROHO territories have been analysed in detail (Fig. 4 and Table S3).

**Fig. 4:**
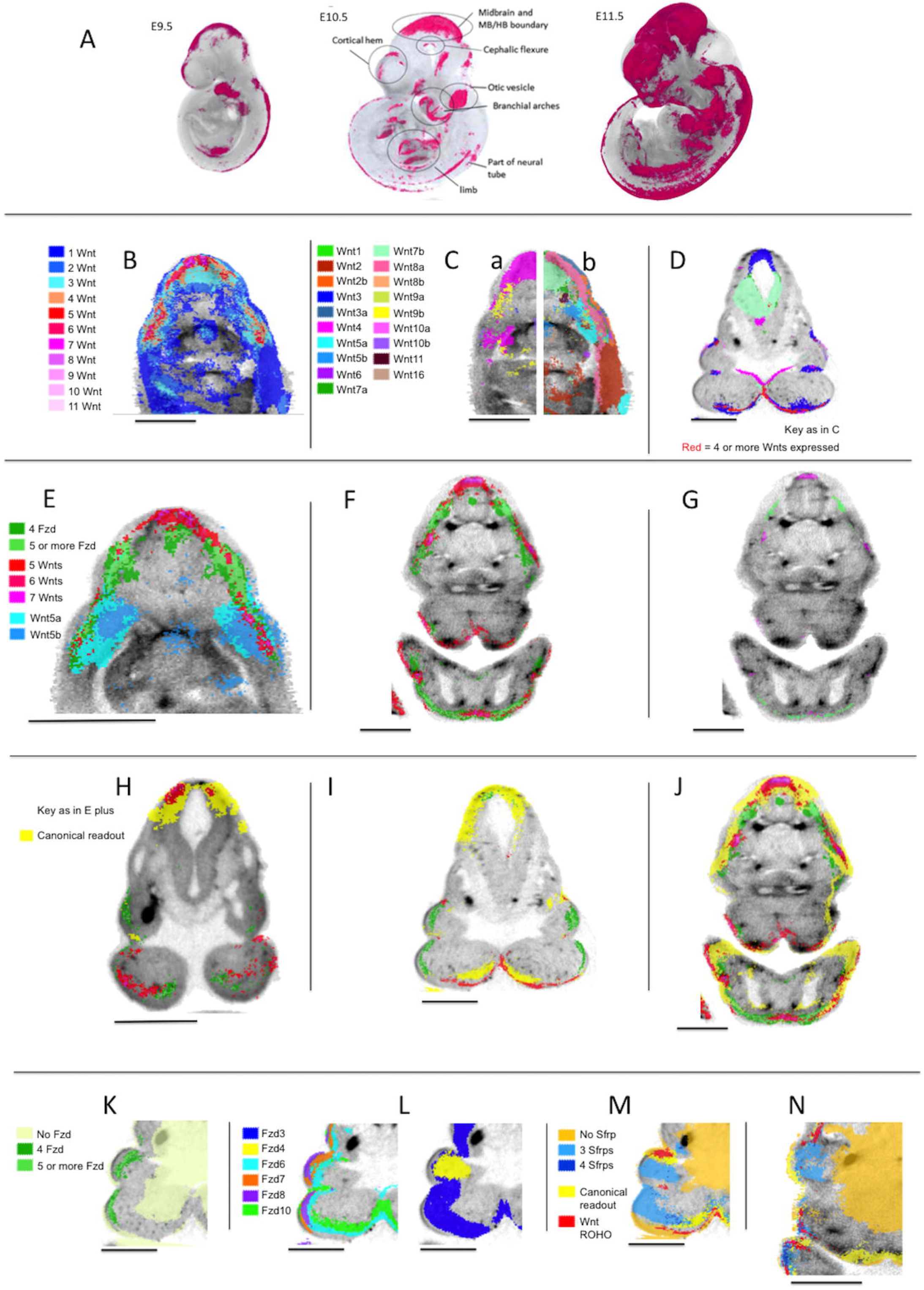
Regions of high occupancy Wnt and Fzd expression. A) Regions of high occupancy of Wnt gene expression (red) at E9.5, E10.5 and E11.5. B) Transverse section through an OPT reference model of an E11.5 embryo in the mid-flank region showing the distribution of occupancy of Wnt expression as indicated. This section reveals three Regions of High Occupancy (ROHOs) C) Same section as B, showing the mapped expression of individual Wnt genes as indicated. For clarity, (a) left and (b) right halves of the section are shown with the expression of (a) Wnts 3, 4, 8b, 9b, 10a, 10b and (b) Wnts 1, 2, 2b, 3a, 5a, 5b, 6, 7a, 7b, 8a, 9a, 11, 16. D) A section through an OPT reference model of an E10.5 embryo in the mandibular region. Regions of high occupancy of Wnt expression (Red; 4 or more Wnts expressed) and individual Wnt gene expression domains are shown for Wnts 3, 4, 7a, 7b, 10a and 10b (key as in C). E) Transverse section through an OPT reference model of an E11.5 embryo in the mid-flank region showing Fzd ROHOs (key as indicated) in the context of Wnt ROHOs (key as in B showing only 5+ Wnt genes) and expression of Wnt5a and Wnt5b. F) Transverse section through an OPT reference model of an E11.5 embryo in the mandibular region showing Wnt and Fzd ROHOs (key as in E). G) The same section as in F, showing only the peaks of Wnt and Fzd occupancy (Wnt occupancy of 7 and more; Fzd occupancy of 6 and more. H, I, J) sections through the mandibular region in OPT reference models of embryos at E9.5, 10.5 and 11.5 respectively. The images show the canonical pathway readout (Tcf/Lef-GFP RNA) (yellow) in the context of Wnt and Fzd ROHOs (key as in E). K) Section through the mandibular region of an OPT reference model of an E10.5 embryo showing Fzd ROHOs compared with where no Fzds are detected (as indicated) L) The same section as in K showing the expression of individual Fzds genes as indicated (the section is repeated for clarity). M) The same section as in K showing Sfrp occupancy 0, 3 and 4 (as indicated) in the context of Wnt ROHOs (key as in E) and canonical pathway readout (Tcf/Lef-GFP) in yellow. N) The same key as N on a section through the maxillary and mandibular region of an OPT reference model of an E11.5 embryo.

In general, each ROHO defined in this way has a consistent location through successive stages (Fig. 4A). The number of Wnt ROHOs is approximately the same at E9.5 and E10.5 and increases at E11.5 when many Wnt ROHOs are more extensive and have higher levels of occupancy (Fig. S4C). Fzd ROHOs also become more extensive between E9.5 and E11.5.

Broadly speaking, Wnt ROHOs and Fzd ROHOs are localised to the same parts of the embryo, but not in all cases (Fig. 4E-I). In particular, many peaks of occupancy in Wnt ROHOs do not coincide with peaks in Fzd ROHOs, even at E11.5 when intersection between Wnt and Fzd ROHOs is maximal (Fig. 4F,G)

#### The expression of individual Wnts in relation to regions of multiple Wnt expression

For most Wnt genes, expression is predominantly in domains that intersect Wnt ROHOs (compare Fig. 4C with 4B). However, the existence of Wnt ROHOs is not simply the result of the random intersection of large, unrelated expression domains. The Wnt genes most commonly expressed in ROHOs typically have small to medium-sized domains that individually intersect discrete ROHOs and may extend through the adjacent epithelium or mesenchyme in a manner that appears to be localised around the ROHO (Fig. 4C,D,E). The ROHO-related expression of, for example, Wnt3 at all stages (Fig. 4D) and Wnt7a at E11.5, is largely epithelial. Others, for example Wnt5a, have expression domains that include the epithelium of the ROHO and extend into sub-adjacent mesenchyme (Fig. 4E).

For a few Wnts, e.g. Wnt3 (Fig. 4D), most expression is ROHO-related. However, the majority have both ROHO-related and apparently non-related expression domains. Wnt2 has extensive expression domains in the ventral trunk that display no consistent relation to ROHOs. Wnt6, though expressed in ROHOs, is expressed quite widely in surface epithelium and does not display convincingly localised expression in ROHOs. However, with the single exception of Wnt16, each Wnt displays at least one instance of local expression in a ROHO at one of the stages we examined.

For a selection of 36 Wnt ROHOs, we examined serial virtual sections to determine which genes have expression domains that intersect any voxel in the ROHO; referred to as the gene set for that ROHO (Table S3 shows ROHO location and gene set; W1-W36, counting ROHOs at each stage as separate). The number of ROHOs in which each Wnt is expressed and the distribution of the number of Wnts per ROHO are shown in Fig. S3A and C. It can be seen in Fig. S3A that Wnt3, Wnt4, Wnt10a *or* Wnt10b are expressed in almost all the ROHOs we examined. In addition, though their expression is more widespread, Wnt7a or Wnt7b, or both, are expressed in 33/36 ROHOs. Thus, Wnts 3, 4, 7a, 7b, 10a and 10b are commonly expressed in Wnt ROHOs across the three stages (see, for example, Fig. 4D). Wnt3a, could arguably be considered as a member of this common Wnt ROHO set at E9.5 when, strikingly, it is expressed locally and almost specifically in ROHOs.

Apart from the common Wnt ROHO gene set described above, the composition of expression in Wnt ROHOs is dynamic. As development proceeds through the stages we examined, there is a general increase in the number of Wnts expressed in ROHOs (Table S3; Fig. S3C). At E11.5, with only 7 exceptions (Wnt2/2b in W16; Wnt5a/5b in W19; Wnt8a/8b in W13, W14, W21; Wnt9a/9b in W19, W20), each paralogous pair is represented by at least one member in each of the 14 ROHOs examined at that stage. There are some notable instances, for example, Wnt10a and Wnt10b, where the expression of a paralogue at one stage is apparently substituted by its partner at the following stage (Table S3).

#### The expression of individual Fzds in relation to regions of multiple Fzd expression

Like the Wnts, most Fzd expression domains intersect Fzd ROHOs examples are Fzds 3, 6, 7, 8 and 10 (Fig. 4L). Unlike Wnt ROHOs, Fzd ROHOs are characterized by the expression of most members of the gene family (Table S3; Fig. S3B). Five of the ten Fzds (3, 6, 7, 8, 10) are expressed in all or nearly all Fzd ROHOs, two (Fzds 1 and 9) are expressed in more than half the Fzd ROHOs we examined and two (Fzds 4 and 5) are expressed in about one third of Fzd ROHOs. 31 of the 39 Fzd ROHOs we examined express 6 or more Fzd genes (Fig. S3D).

#### Wnt and Fzd ROHOs and canonical Wnt pathway activity

At all three stages, most Wnt ROHOs and Fzd ROHOs show at least partial intersection with Tcf/Lef-GFP activity. In some instances, the correlation is tight (for example the Wnt ROHOs in the distal forelimb and distal hindlimb at E10.5), but in the majority of cases Tcf/Lef-GFP activity, either partly or wholly intersecting the ROHO, extends beyond the ROHO (Fig. 4H,I,J). Some discrete domains of Tcf/Lef-GFP activity, though not intersecting any ROHO, lie in tissue immediately adjacent to one. One example is the isthmus (Table S3: W17, F17) where Tcf/Lef-GFP activity is absent in the flexure but present in the adjacent neural tissue. There are a few instances where Wnt and Fzd ROHOs do not display any apparent correlation with Tcf/Lef-GFP activity, for example in the mandible (Table S3: W2 and F2) at E9.5 (Fig. 4H). However, in these cases, Tcf/Lef-GFP is active at the subsequent stage, intersecting with the corresponding ROHO (Fig. 4I).

### Comparison of patterns: similarity and divergence of paralogous pairs of Wnt genes

We previously compared the expression of four pairs of Wnt paralogues (Wnt2, Wnt5, Wnt7 and Wnt8) between mouse and chick embryos (Martin et al., 2012). Here we compare seven pairs of paralogues (Wnt1, 3, 5, 7, 8, 9, 10) for overall similarity and divergence of expression in the mouse. Fig. 5 compares the whole embryo expression patterns at E10.5 and Table 1 shows the Jaccard similarity indices (JI) for each pair in the whole embryo (Jaccard Index = volume of intersection/volume of union of the two domains). Wnt7a and Wnt7b show the greatest similarity of any pair of Wnt genes across all stages with extensive overlap in the CNS (Table 1; Fig. 5D; Table S4).

**Fig. 5:**
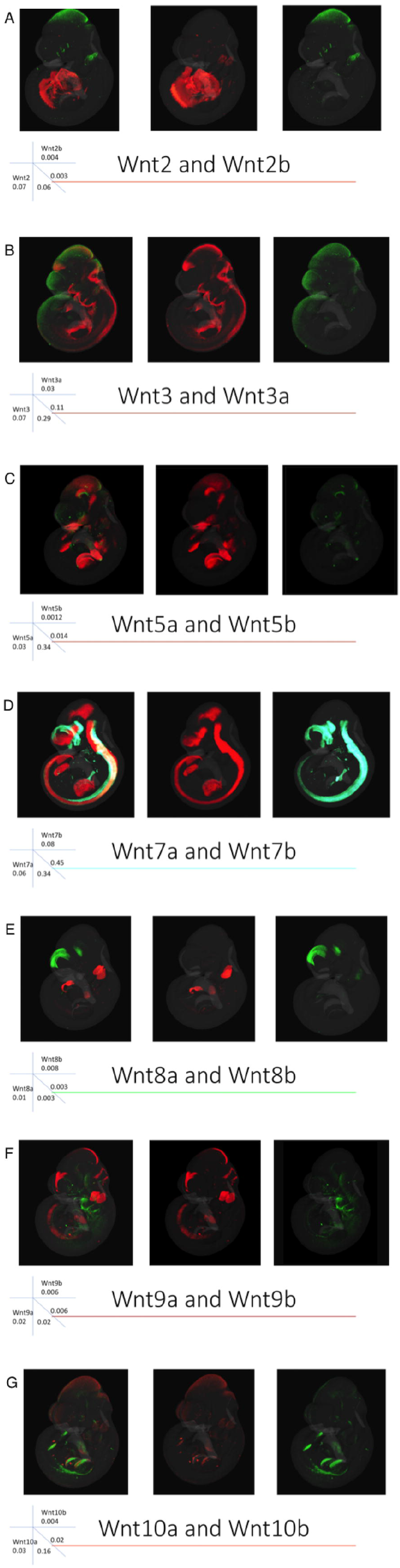
Comparison of expression of paralogous pairs of Wnt genes at E10.5 mapped to the reference embryo model. Each row shows a different pair of the seven Wnt paralogues, as indicated. The combined image of both genes is shown on the left and the two individual patterns in the order listed from left to right. The rubric indicates the relative size of each domain (e.g. Wnt2 occupies 7% of the embryo); the intersecting numbers show the proportion of one pattern overlapping the other so 6% of the Wnt2b domain overlaps the Wnt2 domain. Note the highest level of overlap for the Wnt7 paralogues, followed by the Wnt3 and Wnt5 paralogues.

**Table 1:**
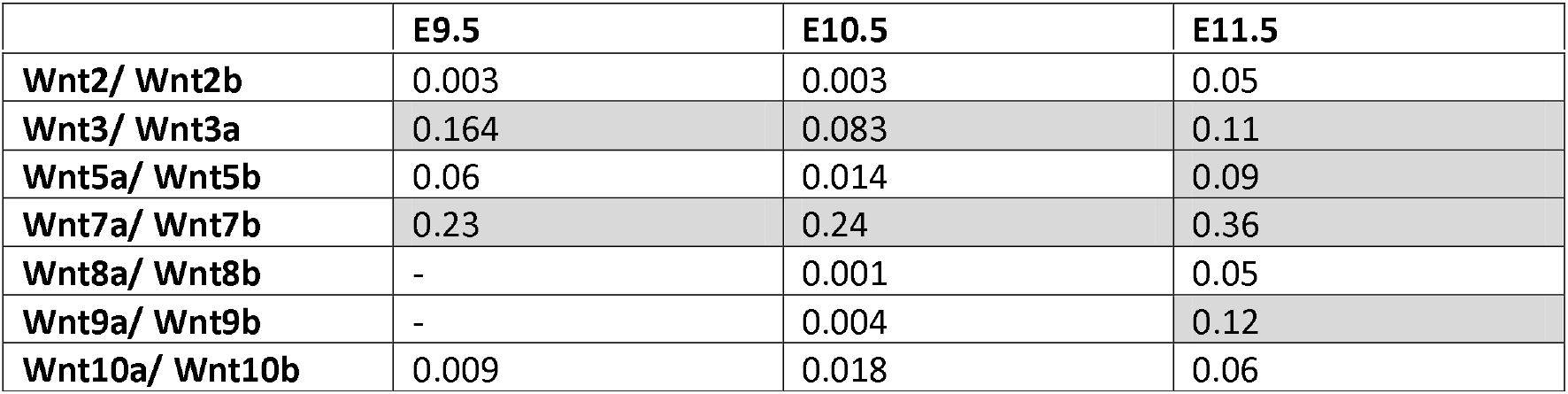
Similarity indices (Jaccard (JI)) of expression domains of Wnt paralogous gene pairs over time (highest values are highlighted in grey)

|  | E9.5 | E10.5 | E11.5 |
| --- | --- | --- | --- |
| Wnt2/ Wnt2b | 0.003 | 0.003 | 0.05 |
| Wnt3/ Wnt3a | 0.164 | 0.083 | 0.11 |
| Wnt5a/ Wnt5b | 0.06 | 0.014 | 0.09 |
| Wnt7a/ Wnt7b | 0.23 | 0.24 | 0.36 |
| Wnt8a/ Wnt8b | - | 0.001 | 0.05 |
| Wnt9a/ Wnt9b | - | 0.004 | 0.12 |
| Wnt10a/ Wnt10b | 0.009 | 0.018 | 0.06 |

Their expression patterns however show complementarity especially in the dorsal and ventral aspects of the forebrain (Fig. 5D). Wnt7a is more widely expressed than Wnt7b in the neural tube and limb. The Wnt3 paralogues also show extensive similarity with the second highest JI at E9.5 among any Wnt gene pair and a high score across stages. Although distinct, extensive overlap in the midbrain is clear (Fig. 5B). There is also overlap in the posterior neural tube. The Wnt10 paralogues are distinct at E9.5 (Wnt10b expression is extensive at this stage and more similar to other patterns) and most similar to each other at E10.5 and E11.5, particularly in the limbs, although the level of similarity is low for E11.5 (0.06). Wnt5 and Wnt9 paralogues show increased expression similarity over time, while Wnt5 genes become more divergent from other patterns in general. Wnt2 and Wnt8 paralogues show most distinct patterns with weaker similarity indices than most other pairs of non-paralogous Wnt genes (Table S4).

### Comparison of patterns: all gene expression patterns and canonical pathway activity

The expression domains of all genes and canonical readout were examined for overlap using parallel-coordinate analysis, JI similarity (Table S4) and visual comparison. Fig. 6A shows striking similarity between where 3+ Wnts are co-expressed and canonical pathway readout, compared to where 0 or unique genes are expressed. Fig. 6A also illustrates each component gene expression pattern at E11.5. Table S4 shows the JI for each pair of genes. Plotting pairwise JI across stages (Fig. 6C), it is clear that Wnt gene family expression patterns generally become more similar to each other over time, with some notable exceptions (Wnt1, Wnt6, and Wnt10). This is also the case when comparing across Wnt and Fzd gene families. For example, while 50% of the Wnt16 expression domain lies outside any Fzd expression domain at E9.5, this drops to 13% at later stages. This is also reflected in the proportion of any pattern that is uniquely expressed (Table S2).

**Fig. 6:**
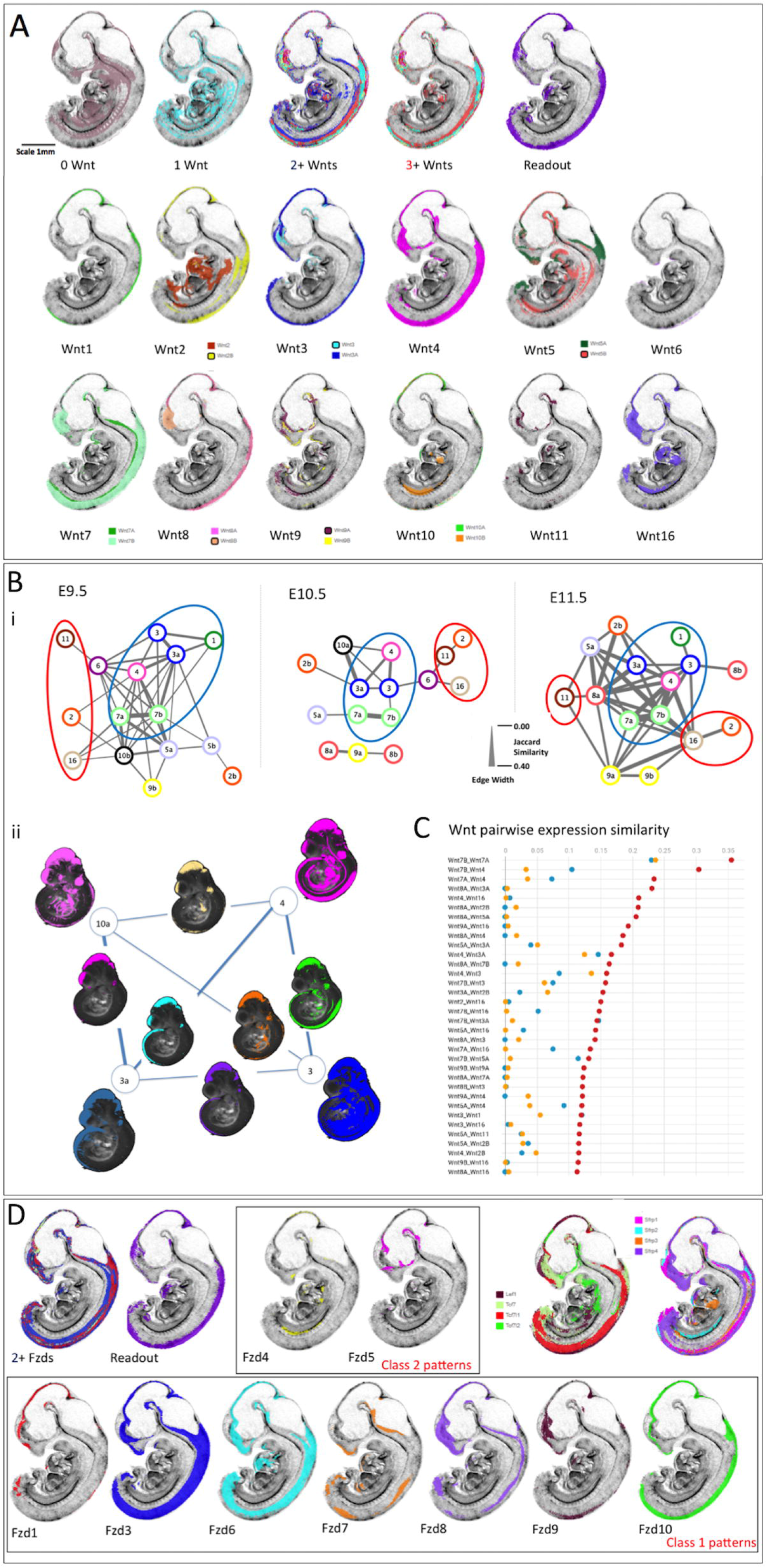
Integrated comparison of gene expression pattern similarity across Wnt, Fzd and other pathway component genes. A) E11.5 example of the visual analysis carried out; all virtual sections are identical, mid-sagittal. The top row shows the territories where zero Wnts are detected (0 Wnt), where individual Wnt genes are expressed uniquely (1 Wnt), where 2 or more (2+ Wnts) or 3 or more (3+ Wnts) genes are co-expressed and where canonical Wnt pathway readout is detected (Tcf/Lef-GFP). Rows 2 and 3 represent mapped expression of each of the Wnt family genes as indicated. The analysis included viewing the full set of sections in all orientations and across stages. Bi) Network diagrams representing the similarities between Wnt expression pattern across time. The lines connecting nodes represent the Jaccard Index (JI) of similarity (intersection/union), with thickness scaled as shown. Each “node” represents a Wnt gene as indicated (e.g. 3a=Wnt3a). For comparison of the network, thresholds were adjusted to show the 15 genes with expression patterns most similar to other Wnt genes at each stage. Blue circles enclose the group with most highly similar expression patterns, consistent across stages (Group 1). Red circles enclose the most divergent expression patterns (Group 3). Bii) illustrates visually the nature of the lines connecting genes in the network focussing on the most highly connected genes at E10.5; Wnts 3, 3a, 4 and 10a. The mapped Wnt expression patterns are shown here in projection through a 3D view of the reference model embryo, at each corner of the network (as indicated). Intersection domains, where each pair of expression patterns intersect, are shown on the lines connecting that gene pair. C) Top 34 similarity scores among all Wnt genes across stages. The horizontal axis refers to Jaccard Index. (red, E11.5; blue E10.5; yellow, E9.5). D) Domains of multiple Fzd expression patterns (2+ Fzds) correspond well to territories of canonical pathway readout (Tcd-Lef-GFP). Fzd expression patterns fall within two classes: class 1 (row 2) are similar to canonical readout, Tcf/Lef transcription factor and Sfrp family member expression patterns (row 1, right). Example patterns at E11.5 are shown.

Wnt gene similarity indices were plotted as networks to compare the relationships between expression patterns across stages. For each stage, we plotted network graphs in which genes are presented as nodes connected by “edges” with a line thickness that represents the similarity (JI) between expression patterns. Fig. 6B illustrates networks for the 15 Wnt genes that display the most similar patterns to other Wnts. The patterns can be divided into 3 groups:

#### Group 1

Core set of genes with expression patterns that show high JIs when paired (Fig.6Bi; blue circled). This includes Wnt7 and Wnt3 paralogues, and Wnt4. Wnt1 is also included at E10.5 and E11.5. These expression patterns are most similar to canonical readout (Fig. 6A; Table S4). Wnt4 changes somewhat over time; while consistently similar to Wnts3, 7 and Wnt5a, it becomes more similar also to other patterns (Wnt8a, Wnt16, Wnt9, Wnt10, Wnt2b) at E11.5, due largely to a new territory of forebrain expression (Fig. 6B).

#### Group 2

Wnt11, Wnt16 and Wnt2 show reduced similarity across stages (Fig. 6Bi; red circled) and are among the genes with the largest proportion of unique expression (Table S2; Fig. 3C). Visual analysis shows the patterns are distinct from canonical read-out (Fig. 6A). Both Wnt11 and Wnt16 show low similarity generally at E9.5 with some similarity to each other by E10.5 (0.04). Both begin to be expressed in the brain at E11.5 driving increased similarity to other patterns at that stage; however, while Wnt16 shares brain expression domains with more commonly expressed Wnt genes it is most similar to Wnt2, Wnt9b and Wnt11 in the trunk.

#### Group 3

The remaining gene expression patterns (Fig. 6Bi, uncircled) show intermediate similarity that can vary across stages; at some stages they may be more similar to Group 1 genes. These include Wnt5, Wnt8, Wnt9 and Wnt10 paralogues, and Wnt2b. It is striking that both Wnt8a and Wnt8b patterns are most similar to Wnt9a at E10.5 although they show very low similarity to each other; Wnt9a shares different aspects of both patterns. Wnts 9a and 9b become more similar to each other and share expression territories most with Wnt16 at E11.5. Wnt2b becomes more typical of Group 1 patterns at E11.5. At E9.5 Wnt5a shows strong similarity to Wnt7 paralogues while Wnt5b shows an intermediate pattern; however, Wnt5b expression becomes more distinct from other Wnt patterns with time; at E11.5 it shows some similarity with Group 1 genes largely through neural expression while it intersects Wnt16 and Wnt2 in the trunk. Wnt10b expression is extensive at E9.5 driving more similarity with Group 1 genes, but the pattern is overall very distinct. At E10.5 Wnt10a becomes more similar to Group 1, largely due to midbrain expression while Wnt10b is more similar to Wnt2 and Wnt16.

The expression pattern of each Wnt gene generally comprises several sub-domains that are shared with some other Wnts but absent or much reduced in others. Thus, for example, at E10.5, Wnt expression in the dorso-lateral neuro-epithelium of the future telencephalon, in the anterior neural tube, mandible, branchial arches (grooves and pouches) and proximal limb comprises different combinations of genes. We examined the pattern similarities represented by the lines connecting pairs of genes in the network graph (Fig.6Bi (edges)) from this perspective. Fig. 6Bii presents a visual picture of edges in the network graph: in some instances, multiple connections to the same gene in the network reflect similar, though not identical, sets of intersecting expression domains (e.g. Wnt3a to Wnt10a and to Wnt4 at E10.5), whereas, in other instances, different connections to the same gene reflect different combinations of intersecting domains (e.g. Wnt4 to Wnt3a and to Wnt3 at E10.5). Fig. 6Bii strikingly reveals that a domain centred on the midbrain is shared by all group 1 genes.

Pair-wise comparison reveals that Fzd gene expression falls broadly into two pattern classes. Class 1; Fzd 3, 6, 7, 8, 9, 10, and to a lesser extent Fzd1, show general overlap and similarity of expression across stages (Fzd2 is not detected at any stage, Fzd5 is not detected and Fzd1 very little expression at E9.5; Fzd9 expression is restricted largely to the brain) (Fig. 6D). Class 1 patterns also show general overlap with canonical pathway readout (at E11.5, Fzd versus readout JIs range from 0.26 (Fzd10) to 0.01 (Fzd9)), with extensive commonality with expression of the Tcf/Lef transcription factors and Sfrp modulators. In contrast, Class 2 patterns, Fzd4 and Fzd5, show much less similarity with readout (0.057 for Fzd4 and 0.025 Fzd5), each with distinct patterns. These different groups of Fzd patterns also overlap with different Wnt expression patterns, although the individual Wnts involved are dynamic over time. While Class 1 Fzd patterns predominantly show similarity with Wnt Group 1, Fzd4 and Fzd5 both show greatest similarity to Wnt9a and Wnt8a at E10.5, with increased similarity with Wnt16 (0.124 and 0.149) and Wnt11 (0.059 and 0.12) at E11.5.

Among Tcf-Lef transcription factor patterns, Tcf7 and Lef1 are most similar and Tcf7l2 is the most divergent pattern across stages (Fig. 6D; Table S4). Sfrp gene expression patterns are dynamic, with Sfrp4 more divergent at E10.5 but more similar, particularly to Sfrp1 by E11.5 (change from 0.15 to 0.33 JI) (Fig. 6D) (note Sfrp5 expression was not detected). Sfrp1-4 patterns were also analysed visually for co-expression (Fig. 4M,N). Regions with co-expression of 3 and 4 Sfrp genes are generally well-defined and discrete. At E10.5, these regions rarely coincide with Wnt ROHOs; e.g. in the core mesenchyme and distal epithelium of the developing branchial arches (Fig. 4M). However, by E11.5, domains of multiple Sfrp gene expression intersect the expanding Wnt ROHOs, for example within, or close to the surface epithelium in the developing face and limbs (Fig. 4N).

The Fzds show some interesting relationships to Sfrp patterns, but the Wnts are more complex and do not show consistent general relations with Sfrps. For example, at E10.5, the group of Fzds 6, 7, 8 and 10 show similarities of expression to Sfrp2 in the face while Fzds 4, 5 and 9 patterns display no apparent relation to Sfrp2 expression. At E11.5, Sfrp2 intersects Fzd7 expression in interesting patterns in the face, limb and trunk, while Sfrp4 intersects Fzd7 in an interesting pattern in the limb.

### Detailed analysis of integrated expression patterns in the ventral diencephalon

Using the resource provided here novel insights can be gleaned through focused analysis of any region of the embryo and used to build testable hypotheses. For example, examining the ventral diencephalon (VD) at E10.5 revealed a striking complementarity between Shh and canonical Wnt readout (Fig. 7A-D). Tcf/Lef-GFP showed a gradient of expression through the midline of the VD that was strongest in the peduncular hypothalamus and the terminal hypothalamus caudal to the infundibulum. In particular, 3D rendered images of mapped data show that the Shh expression domain surrounds the Tcf/Lef-GFP domain (Fig. 7C).

**Fig. 7:**
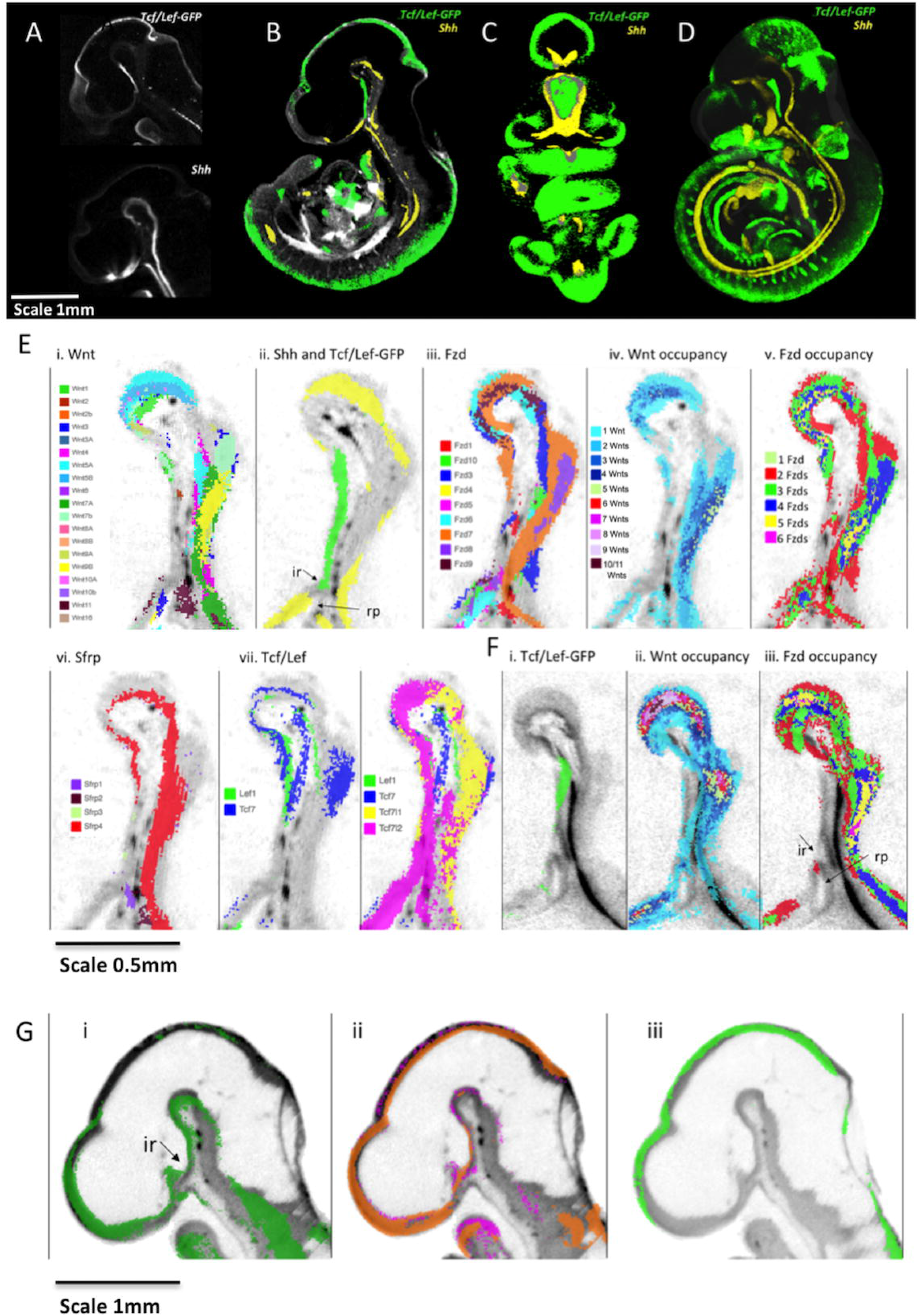
Integrated expression patterns in the Ventral Diencephalon: Complementary expression of Shh and canonical Wnt pathway readout. A) Tcf/Lef-GFP readout pattern and Shh expression in virtual sagittal sections through the diencephalon of raw OPT data. B-D) Mapped data with grayscale render of the 3D model showing anatomy. B shows a virtual sagittal section through the 3D model emphasising the complementary expression of Tcf/Lef-GFP readout pattern (green) and Shh (yellow) in the ventral diencephalon. C is a thick virtual coronal section (288 μm) through the 3D model of mapped data again showing complementarity in the patterns (absence of overlap verified on serial sections) D is a full 3D representation (point cloud render). Scale bar in C and D as for B. E) Sagittal sections through the ventral diencephalon at E10.5 showing: i) mapped expression of all <u>Wnt</u> genes as indicated by the key; ii) canonical Wnt readout (Tcf/Lef-GFP, green) and Shh (yellow) expression on the same section; iii) mapped expression of all Fzd genes as indicated by the key; iv) Wnt occupancy; the number of Wnt genes co-expressed as indicated (no colour indicates zero Wnts are detected); v) Fzd occupancy; the number of Fzd genes co-expressed as indicated (no colour indicates zero genes detected); vi) mapped expression of Sfrp genes as indicated by key; vii) mapped expression of Tcf/Lef transcription factor genes as indicated by key. F) Sagittal sections through the ventral diencephalon at E11.5 showing i) canonical Wnt readout (Tcf/Lef-GFP, green); ii) the number of Wnt genes co-expressed (key as in E iv) and iii) the number of Fzd genes co-expressed (key as in E v) (no colour indicates zero genes detected). G) Sagittal sections through the brain at E9.5 showing i) mapped expression of Wnt7a (green), ii) mapped expression of Fzd5 (magenta) and Fzd7 (orange) and iii) canonical Wnt readout (Tcf/Lef-GFP, green). Abbreviations: *ir* infundibular recess; *rp* Rathke’s pouch

We asked what Wnt and Fzd expression combinations might drive expression of the Tcf/Lef-GFP reporter in the VD by digitally segmenting (Baldock et al., 2003; Brune et al., 1999) the VD anatomical domain in the E10.5 reference model and examining the mapped expression of genes in the Wnt signalling system using parallel co-ordinate analysis. We asked which genes are detected a) where Tcf/Lef-GFP is active and b) where Shh is expressed (Fig. S4). We then confirmed expression of these Wnt and Fzd genes in the VD neuroepithelium and Rathke’s pouch through visualisation of mid-sagittal sections of both mapped and original 3D OPT data. Visual examination revealed that the region positive for canonical Wnt-readout shows limited detection of Wnt and Fzd expression at E10.5 (Fig. 7Ei-iii). Indeed, most of the region shows zero Wnt detection and there are only restricted regions of single Wnt genes; Wnt2, Wnt4, Wnt7b and Wnt1 within the territory (Fig. 7Ei). Similarly, much of the region shows no detectable expression of Fzd genes; only Fzd7, Fzd3 and Fzd1 are expressed in restricted sub-regions (Fig. 7Eiii). In contrast, rostrally and caudally, where Shh is detected, multiple Wnts and Fzds are expressed (Fig. 7Eiv and v). For other members of the Wnt signalling system only Sfrp4 is expressed in the cephalic flexure and in the caudal VD but not throughout the rostral VD where the pathway is active (Fig. 7E vi). Among Tcf/Lef transcription factors, there is extensive expression of Tcf7l1 and Tcf7l2 whereas Lef1 and Tcf7 are restricted to the caudal VD (Fig. 7Evii). Ror2 is expressed, most strongly in the caudal VD (Fig. S4). In summary, canonical readout is seen in the VD where Tcf/Lef transcription factors are expressed and Sfrp expression is restricted, but where, in a substantial part of the region, no Wnt or Fzd expression was detected. Shh is expressed where no canonical output is detected.

A day later, at E11.5, the region still has limited Wnt and Fzd gene expression while canonical readout is restricted more caudally (Fig. 7F). To investigate if the activity at E10.5 could be due to earlier expression of Wnts and Fzds we examined data at E9.5 which showed a single Wnt gene, Wnt7A, and two Fzd genes (5 and 7) expressed (Fig. 7G). While we detected no canonical activity at E9.5, these pathway components could be involved in triggering later activity.

## DISCUSSION

Wnt sinalling is one of the most studied sets of biological pathways, yet the challenge to understand the basis of spatial and temporal control of its biological outputs during development remains. Decades of work has described the expression of individual or small sets of genes in the vertebrate system, but the picture that emerges is patchy and driven by the focus of diverse studies. Here we have used the Mouse Atlas approach to capture comprehensively the expression patterns of all Wnt genes, their Fzd receptors, Sfrp modulators, Tcf/Lef transcription factors as well as other interacting factors in mouse embryos in 3D over the developmental period when the patterning of different organ rudiments is being elaborated. Mapping the expression of different genes to common 3D digital models of embryos at each stage enabled an integrated spatio-temporal analysis of the patterns and comparison to a reporter (Ferrer-Vaquer et al., 2010) that reveals activation of one of several pathway outputs, the canonical pathway. This study provides insight into a level of organisation of the patterns not previously apparent, as well as presenting a resource that can be utilised further by the research community.

Our comprehensive approach indicated that the territories where no Wnt and no Fzd expression is detected across the three stages are largely ventral and visceral. Territories where the expression of only one Wnt gene was detected are also predominantly ventral, in particular expression of Wnt2 across stages and Wnt11 at E10.5 (Fig. 3C). Both genes show very divergent expression patterns in our cross-pattern analysis (Fig. 6B) pointing to possible unique roles and regulation. They show mutant phenotypes in the placenta, kidneys and lungs consistent with these patterns (Goss et al., 2009; Majumdar et al., 2003; Monkley et al., 1996). By contrast those domains with expression of multiple genes in all families are in dorsal and lateral regions, particularly the central nervous system, limbs, flank and face, consistent with the primary body axis being determined along the dorsal midline (for example (Arraf et al., 2016). To address the question why there are so many Wnt and Fzd genes in a single organism, we have examined the hypothesis that Wnt and Fzd expression is a mosaic of domains, each expressing only one or a few members of these families. Our results are not consistent with this hypothesis. Some anatomical regions show the expression of one or a few Wnts co-expressed, similar to what would be expected from this hypothesis. But there are many regions co-expressing strikingly large numbers of Wnts and Fzds. The focus of the question thus shifts to why are so many Wnts and Fzds co-expressed in these regions?

A striking finding is the co-expression of a large fraction of all Wnt or Fzd genes in localised regions of the embryo (termed ROHOs). These regions may reflect regulatory “hot spots” for the gene families. Some coincide with known Wnt signalling centres such as the isthmus and the cortical hem but others are novel and have not been previously detected, for example the flank anterior to the forelimb (W5 in Table S4) and the ventral aspect where the forelimb meets the flank (W8 in Table S4). It is important to note that generally each gene has a distinctive expression pattern extending beyond the ROHO, often including domains unconnected to ROHOs. The intersection of patterns in ROHOs generally suggests spatio-temporal regulation centred on a small region of tissue. These observations open an avenue to investigating the signalling characteristics of these regions and their importance in patterning.

Asking how ROHOs relate to canonical pathway activity, there is generally correspondence between Wnt and Fzd ROHOs and pathway readout, but this is neither universal nor precise. The respective peaks of Wnt and Fzd ROHOs are usually offset. The same is true of regions of Tcf/Lef-GFP reporter activity. This suggests that it is not simply the additive effect of multiple Wnt genes that activates the pathway; the relationship is more complex reflecting the full regulatory landscape. Indeed, canonical signalling is not restricted to Wnt and Fzd ROHOs. We have quantified the extent of each expression domain and compared the territories where different numbers of Wnt genes are co-expressed with canonical Wnt readout showing that 33% of readout falls within regions of unique Wnt gene expression at E9.5 and E10.5. We also found canonical Wnt pathway readout in the absence of any detectable concurrent Wnt or Fzd gene expression (e.g. in nasal epithelium at E11.5 and the ventral diencephalon at E10.5 and E11.5). In the case of the VD, earlier (E9.5) expression of individual Wnt and Fzd genes could account for the later activity.

Another striking feature of all Wnt patterns is that they generally become more similar over the developmental time period covered (Fig. 6C); also reflected in the increased overlap of genes in ROHOs over time. Pairwise comparisons of similarity between expression patterns using the Jaccard Index (Table S4) provides a view of the deployment of Wnt function that complements the co-expression analysis. Network analysis of pairwise comparisons (Fig. 6B) revealed three groupings of patterns. The first (Group 1) are similar across stages and often associated with regions of canonical pathway activity and Wnt ROHOs (Table S4; expression of Wnts 3, 7 and 4 occurs in >30/36 ROHOs analysed) while the second (Group 2) are less associated with canonical activity and include the genes most expressed in unique territories and in ventral and visceral domains (Wnts 2 and 11). A third group are intermediate and/or change their similarities over time; e.g. Wnt10a is very closely related to Group 1 patterns at E10.5 but dramatically less so at other stages. Group 1 expression patterns might lie closer to an ancestral pattern aligned with the primary body axis, while other patterns are more divergent and associated with recently added Wnt system functions: e.g. Wnt2 association with placental development and Wnt11 with kidney development. The predominant elements in the similarity between group 1 patterns lie within the CNS including the dorsal midbrain, the cortical hem in the forebrain and the dorsal neural tube. Interestingly, further analysis of the similarity between Group 1 patterns (Fig. 6Bii) suggests that the expression of each of these genes comprises combinations of subdomains shared with some but not all members of the set indicative of modular regulation.

Turning to the question why so many Wnts and Fzds are co-expressed in localised regions, there are three non-exclusive possibilities:

A. Convergent evolution of independent family members to satisfy a functional requirement for the expression of multiple genes, e.g. a threshold for Wnt ligand concentration.
B. Conserved regulation of Wnts and Fzds, either active or passive.
C. The existence of spatio-temporal regulatory control that spans the family, i.e. a form of “meta-regulation” of the family. This raises the possibility that there exists a level of regulation, hitherto unknown, that directs the expression of the Wnts as a suite, and similarly for Fzds. One possibility, for example, would be positive feedback regulation across the gene family.

We envisage that these possibilities apply not to the entire expression pattern of any gene, but rather to independently regulated sub-domains of expression. Interestingly, Wnt expression studies in Amphioxus, which has 13 Wnt genes, also shows regions where multiple Wnts are expressed, for example posterior nested expression domains (Somorjai et al., 2018). This suggests that spatio-temporal intersection is not unique to the more complex gene family in the mouse. Evolutionary studies comparing Amphioxus and the tunicate *Oikopleura dioica* have suggested three modes of evolutionary change in the Wnt gene family namely, conservation of function, function shuffling and gene loss (Martí-Solans et al., 2021). It is possible that the co-expression of subdomains of different Wnts in the mouse reflects the operation of a conserved ancestral regulation, though not necessarily conservation of precise gene function, across different Wnts.

We previously compared the expression of four pairs of Wnt paralogues within and between mouse and chick embryos (Wnt2, Wnt5, Wnt7 and Wnt8) (Martin et al., 2012) showing evidence of greater divergence between subgroup paralogues than the respective orthologues, consistent with conserved subfunctionalisation/neofunctionalisation in the common vertebrate ancestor. Here we compare all seven paralogue gene pairs reinforcing earlier observations and adding new insight. The Jaccard Index shows that Wnt7 and Wnt3 paralogues are most similar of all Wnt pairwise patterns (Table 1 and Table S4), yet the patterns are distinct, often complimentary, in the same anatomical region. Contrastingly, Wnt2 and Wnt8 paralogues have diverged enormously in their expression characteristics, across all stages. Wnt10 genes present an interesting case where they appear to “swap” territories over time; especially evident in ROHO analysis where the same ROHO switches between expressing Wnt10a and Wnt10b (Table S3). These results add to our understanding of how paralogues arising by duplication of highly conserved genes evolve individually, sometimes maintaining aspects of their regulatory inputs while adjusting precise expression domains within that territory, and/or by acquiring new territories of expression. These finding present interesting contrasting cases (e.g. Wnt7 vs Wnt2 respectively) to dissect the regulatory inputs for each gene pair to fully understand the regulatory changes involved.

In addition to the global analysis described above, our results can be used with a focus on individual organs and as a resource to complement hypothesis-driven approaches. As a case study, we analysed the ventral diencephalon (VD) in some detail. The VD goes on to form the hypothalamus and the neurohypophysis that innervates the oral ectoderm-derived pituitary (adenohypophysis) through the infundibular stalk; together these components form the hypothalamic-pituitary axis, of major importance in homeostasis. Initially using Shh as a marker gene, we immediately noticed a striking complimentary pattern between Shh and the Tcf/Lef-GFP reporter (Fig. 7), suggestive of a repressive relationship between canonical Wnt signalling and Shh expression in this territory. Indeed Osmundsen et al. (Camper et al., 2017; Osmundsen et al., 2017) have demonstrated that rostral expansion of beta-catenin activity leads to coincident loss of Shh expression, elegantly demonstrating our original conjecture from visual analysis.

We further suggest a role for Wnt/Beta-catenin signalling in development of the neurohypophysis, in particular the evaginating infundibulum, and reveal Wnt signalling pathway gene expression patterns that could contribute to this important regulatory output. By digitally dissecting the territories that express Shh and those which show Tcf/Lef-GFP activity, coupled with parallel-coordinates visualisation, we could find potential regulatory inputs to the observed Tcf/Lef-GFP output; i.e. the cocktail of Wnts, Fzds and other regulatory components that are expressed in the region. Surprisingly few Wnts and Fzds are expressed in the region of Tcf/Lef-GFP activity, and none throughout the region at E10.5 and 11.5 whereas very many genes are expressed in the Shh +ve territory where Tcf/Lef-GFP is not active. However, at the earlier stage of E9.5 Wnt7a and Fzd5 and 7 are expressed in the region that later becomes Tcf/Lef-GFP. This cautions against drawing conclusions about pathway activity based on component gene expression patterns alone.

To further explore the ventral diencephalon, making use of the Mouse Atlas EMAGE database, we carried out a spatial query for genes with similar patterns to Shh and Tcf/Lef-GFP. This identified *Vax1* with a complementary pattern to Tcf/Lef-GFP. Vax genes are of particular interest since they are known to inhibit canonical Wnt signalling through activation of an internal promoter transcribing a dominant-negative isoform of Tcf7l2 (Vacik et al., 2011). Furthermore they are dependent on Shh signalling (Zhao et al., 2010) consistent with a mutually repressive relationship between Shh and Wnt signalling through expression of Vax1. We hypothesise that this Shh-dependent inhibition of Wnt/Beta-catenin signalling in the rostral VD is necessary to limit the pituitary forming territory, consistent with ectopic pituitary formation in Vax1-deficient mice (Bharti et al., 2011)

The data reported here can be used to help direct future investigation of the global regulation and function of Wnt and Fzd family genes. In particular, it will be interesting to investigate the effects of manipulating individual genes on the expression and function of co-expressed members of the family. By providing a means to directly visualise comparisons between data in 3D and to incorporate retrospective and future data, the approach provides an opportunity to complement the data reported here with mutational and high resolution, multiplex approaches (Lohoff et al., 2021).

## EXPERIMENTAL PROCEDURES

Expression patterns were generated by *in situ* hybridisation as previously described (Summerhurst et al., 2008) for Sfrp1-4, Ror2, Wif1, and Wise, including previously reported Wnts, Fzds and Tcf/Lef genes. The cDNA sequences used for generation of RNA probes are detailed in Supplementary Table 5. Readout of the canonical pathway was revealed using GFP expression (both RNA in situ and anti-GFP immunofluorescence [Invitrogen A11122 1:200]) in a previously characterised transgenic mouse line (Ferrer-Vaquer et al., 2010).

3D imaging was carried out using Optical Projection Tomography (OPT) as previously described (Summerhurst et al., 2008).

Mouse embryos are referred to by embryonic day (E), however, embryos analysed were staged according to Theiler criteria (Theiler, 1989) to Stages 15, 17 and 19, referred as E9.5, E10.5 and E11.5 respectively. Expression patterns are submitted to the Edinburgh Mouse Atlas of Gene Expression (EMAGE; IDs noted on Table S5), available at https://www.emouseatlas.org/emage/home.php. Patterns can also be viewed openly at https://www.tcd.ie/Zoology/research/groups/murphy/WntPathway/

### Mapping of gene expression data

3D gene and reporter patterns were mapped onto reference embryos at each stage using a manual image-editing tool WlzWarp (Hill et al., 2022). This uses *Constrained Distance Transform* (Hill and Baldock, 2015) which can deliver the complex non-linear transforms required for the variable shape and pose of mouse embryos. WlzWarp is an open-source tool (github.com/ma-tech) and provides interactive non-linear spatial mapping of 3D image data. This has been used for mapping significant volumes of gene-expression data and tested by mapping multiple images of the same gene from independent samples (Hill et al., 2022). Fig. 1 illustrates mapping examples. Mapping accuracy was assessed for each gene by comparing virtual sections of original 3D data (unmapped) against the mapped data (Fig. S1) showing good fidelity in most cases, reduced when mapping surface (ectodermal) expression (Fig. S1F).

### Analysis of integrated data

Primary checking and visual analysis of 3D expression data and mapped patterns used open-source tools MAPaint and MA3DView (github.com/ma-tech). For visualisation of mapped patterns we used the IIP Viewer providing access to the data within a standard web-browser (Armit et al., 2015; Husz et al., 2012). The IIPViewer allows the user interactive selection of arbitrary section views through the mouse embryo and an overlay of all or any combination of the gene-expression patterns (available at www.emouseatlas.org/WntAnalysis). For convenience we have included many of the derived patterns of multiple gene-occupancy including regions where a single gene within a gene family is expressed.

In addition to this section-based visualization we provide a full-3D rendered view using the “point-cloud” approach which delivers a volumetric style view of the entire pattern. Again, viewing any gene-combination can be interactively selected including the entire gene-set. The IIPViewer and point-cloud software and tools for generating the associated data are all open-source from the GitHub ma-tech repositories.

Analysis of mapped expression regions was undertaken using bespoke software tools based on the Woolz image-processing system (Piper and Rutovitz, 1985); github.com/ma-tech/woolz). The tools are csh-scripts that can be executed on any Unix-based system (e.g. Linux, Mac OSX) and generate all of the data values used for the downstream analysis. Specifically they generate:

1. Tables of pair-wise intersection volumes as a count of the number of voxels in common between the two patterns normalised either by the test-pattern volume (row-normalised) or the target pattern volume (column normalised). The overall volumes are provided to enable calculation of absolute volume values and using the voxel resolution these can be converted to real-space (μm^3^) values;
2. Tables of pair-wise similarity values using the Jaccard index (Levandowsky and Winter, 1971) based on the voxel set intersection and union volumes;
3. Volumetric domains of gene-occupancy which for a given gene set (e.g. Wnt) are calculated from the gene-count, i.e. number of genes expressed at every voxel location within the embryo. This occupancy “image” is then thresholded to define regions where for example there are 5 or more Wnts expressed at the same location. This can then be further analysed to reveal which genes are expressed within that region. Such occupancy data was used to reveal the regions of high-occupancy (ROHO) as well as regions of single gene occupancy where there is no overlapping expression within the gene-family;
4. Re-formatted data for visualisation using the IIPViewer, point-cloud viewer and for the parallel-coordinate visual analysis using D3.js (d3js.org) Javascript visualisation library;
5. Re-formatted data for network analysis and input to CytoScape.

All data required for these views are provided in a series of datasets held at the University of Edinburgh public data repository for Wnt Pathway Analysis (Murphy et al., 2021a; Murphy et al., 2021b) https://doi.org/10.7488/ds/3141; https://doi.org/10.7488/ds/3142). In addition, links to the parallel-coordinate views we have used are available at www.emouseatlas.org/WntAnalysis for convenience.

The network analysis software *igraph* (Csardi G and Nepusz T, 2006) igraph.org/, was used to construct networks according to the Jaccard indices of similarity across a variety of threshold levels across stages. The threshold level that showed the top 15 most similar genes at each stage was selected for detailed comparison and network visualisation using cytoscape (Shannon et al., 2003); cytoscape.org) with the network layout unchanged.

## Acknowledgements

We thank Peter Hohenstein and Anna Thornburn for providing Tcf-Lef-GFP embryos used in this study. We thank Yiwen Sun or contribution to mapping of the E9.5 stage. Several students contributed to testing analysis of mapped data, especially Daniel Darby, Jessica Maddock and Cliodhna Smyth. Martina Redmond and Somantha Killion-Connolly assisted with generation of gene expression data. This work was supported by Science Foundation Ireland (Programme Award 02/IN1/B267) and the Irish Research Council.

**Supplementary data Table S1:** Proportional Wnt expression domain volumes across stages (as presented graphically in Fig. 2B).

**Supplementary data Table S2:**

(i) Proportion of the single-Wnt gene expression domain occupied by each Wnt across stages

(ii) (ii) Proportions of each Wnt gene expression domain (and canonical read-out) within the single Wnt expression domain (normalised by the individual Wnt expression domain volume).

**Supplementary data Table S3: Genes expressed in each of the Wnt and Fzd ROHOs we examined.**

**Supplementary data Table S4: Jaccard similarity indices across all gene expression domains and canonical read-out**

**Supplementary data Table 5: details of gene expression probes and EMAGE entry IDs**

## References

Armit, C., Richardson, L., Hill, B., Yang, Y. and Baldock, R. A. (2015). eMouseAtlas informatics: embryo atlas and gene expression database. Mamm Genome 26, 431–440.

Armit, C., Richardson, L., Venkataraman, S., Graham, L., Burton, N., Hill, B., Yang, Y. and Baldock, R. A. (2017). eMouseAtlas: An atlas-based resource for understanding mammalian embryogenesis. Dev Biol 423, 1–11.

Arraf, A. A., Yelin, R., Reshef, I., Kispert, A. and Schultheiss, T. M. (2016). Establishment of the Visceral Embryonic Midline Is a Dynamic Process that Requires Bilaterally Symmetric BMP Signaling. Dev Cell 37, 571–580.

Baldock, R. A., Bard, J. B., Burger, A., Burton, N., Christiansen, J., Feng, G., Hill, B., Houghton, D., Kaufman, M., Rao, J., et al. (2003). EMAP and EMAGE: a framework for understanding spatially organized data. Neuroinformatics 1, 309–325.

Bharti, K., Gasper, M., Bertuzzi, S. and Arnheiter, H. (2011). Lack of the ventral anterior homeodomain transcription factor VAX1 leads to induction of a second pituitary. Development 138, 873–878.

Brune, R. M., Bard, J. B., Dubreuil, C., Guest, E., Hill, W., Kaufman, M., Stark, M., Davidson, D. and Baldock, R. A. (1999). A three-dimensional model of the mouse at embryonic day 9. Dev Biol 216, 457–468.

Camper, S. A., Daly, A. Z., Stallings, C. E. and Ellsworth, B. S. (2017). Hypothalamic β-Catenin Is Essential for FGF8-Mediated Anterior Pituitary Growth: Links to Human Disease. Endocrinology 158, 3322–3324.

Clevers, H., Loh, K. M. and Nusse, R. (2014). Stem cell signaling. An integral program for tissue renewal and regeneration: Wnt signaling and stem cell control. Science 346, 1248012.

Csardi G and Nepusz T (2006). The igraph software package for complex network research. InterJournal Complex Systems 1695.

Davidson, D. and Baldock, R. (2001). Bioinformatics beyond sequence: mapping gene function in the embryo. Nat Rev Genet 2, 409–417.

Ferrer-Vaquer, A., Piliszek, A., Tian, G., Aho, R. J., Dufort, D. and Hadjantonakis, A. K. (2010). A sensitive and bright single-cell resolution live imaging reporter of Wnt/ß-catenin signaling in the mouse. BMC Dev Biol 10, 121.

Goss, A. M., Tian, Y., Tsukiyama, T., Cohen, E. D., Zhou, D., Lu, M. M., Yamaguchi, T. P. and Morrisey, E. E. (2009). Wnt2/2b and beta-catenin signaling are necessary and sufficient to specify lung progenitors in the foregut. Dev Cell 17, 290–298.

Hill, B. and Baldock, R. A. (2015). Constrained distance transforms for spatial atlas registration. BMC Bioinformatics 16, 90.

Hill, B., Husz, Z., Armit, C., Davidson, D. R., Murphy, P. and Baldock, R. A. (2022). WlzWarp: An Open-Source Tool for Complex Alignment of Spatial Data. bioRxiv, 2022.2002.2011.480105. https://doi.org/10.1101/2022.02.11.480105

Holstein, T. W. (2012). The evolution of the Wnt pathway. Cold Spring Harb Perspect Biol 4, a007922.

Husz, Z. L., Burton, N., Hill, B., Milyaev, N. and Baldock, R. A. (2012). Web tools for large-scale 3D biological images and atlases. BMC Bioinformatics 13, 122.

Levandowsky, M. and Winter, D. (1971). Distance between Sets. Nature 234, 34–35.

Loh, K. M., van Amerongen, R. and Nusse, R. (2016). Generating Cellular Diversity and Spatial Form: Wnt Signaling and the Evolution of Multicellular Animals. Dev Cell 38, 643–655.

Lohoff, T., Ghazanfar, S., Missarova, A., Koulena, N., Pierson, N., Griffiths, J. A., Bardot, E. S., Eng, C. L., Tyser, R. C. V., Argelaguet, R., et al. (2021). Integration of spatial and single-cell transcriptomic data elucidates mouse organogenesis. Nat Biotechnol.

Majumdar, A., Vainio, S., Kispert, A., McMahon, J. and McMahon, A. P. (2003). Wnt11 and Ret/Gdnf pathways cooperate in regulating ureteric branching during metanephric kidney development. Development 130, 3175–3185.

Martí-Solans, J., Godoy-Marín, H., Diaz-Gracia, M., Onuma, T. A., Nishida, H., Albalat, R. and Cañestro, C. (2021). Massive Gene Loss and Function Shuffling in Appendicularians Stretch the Boundaries of Chordate Wnt Family Evolution. Front Cell Dev Biol 9, 700827.

Martin, A., Maher, S., Summerhurst, K., Davidson, D. and Murphy, P. (2012). Differential deployment of paralogous Wnt genes in the mouse and chick embryo during development. Evol Dev 14, 178–195.

Monkley, S. J., Delaney, S. J., Pennisi, D. J., Christiansen, J. H. and Wainwright, B. J. (1996). Targeted disruption of the Wnt2 gene results in placentation defects. Development 122, 3343–3353.

Moroz, L. L., Kocot, K. M., Citarella, M. R., Dosung, S., Norekian, T. P., Povolotskaya, I. S., Grigorenko, A. P., Dailey, C., Berezikov, E., Buckley, K. M., et al. (2014). The ctenophore genome and the evolutionary origins of neural systems. Nature 510, 109–114.

Moustafa, R. E. (2011). Parallel coordinate and parallel coordinate density plots. WIREs COMPUTATIONAL STATISTICS 3, 134–148.

Murphy, P., Summerhurst, K., Brady, G., Vendrell, V., Frankel, P., Redmond, M., Lillion-Connolly, S., Armit, C., Hill, B., Venkatarman, S., et al. (2021a). Wnt Pathway Analysis mapped gene-expression point-cloud data. Edinburgh DataShare, 2021.2009.2024. https://doi.org/10.7488/ds/3141

Murphy, P., Summerhurst, K., Brady, G., Vendrell, V., Frankel, P., Redmond, M., Maddock, J., Lillion-Connolly, S., Armit, C., Hill, B., et al. (2021b). Wnt Pathway Analysis mapped gene-expression data. Edinburgh DataShare, 2022.2011.2001. https://doi.org/10.7488/ds/3142

Noelanders, R. and Vleminckx, K. (2017). How Wnt Signaling Builds the Brain: Bridging Development and Disease. Neuroscientist 23, 314–329.

Osmundsen, A. M., Keisler, J. L., Taketo, M. M. and Davis, S. W. (2017). Canonical WNT Signaling Regulates the Pituitary Organizer and Pituitary Gland Formation. Endocrinology 158, 3339–3353.

Piper, J. and Rutovitz, D. (1985). Data structures for image processing in a C language and Unix environment. Pattern Recognit. Lett. 3, 119–129.

Ryan, J. F., Pang, K., Schnitzler, C. E., Nguyen, A. D., Moreland, R. T., Simmons, D. K., Koch, B. J., Francis, W. R., Havlak, P., Smith, S. A., et al. (2013). The genome of the ctenophore Mnemiopsis leidyi and its implications for cell type evolution. Science 342, 1242592.

Shannon, P., Markiel, A., Ozier, O., Baliga, N. S., Wang, J. T., Ramage, D., Amin, N., Schwikowski, B. and Ideker, T. (2003). Cytoscape: a software environment for integrated models of biomolecular interaction networks. Genome Res 13, 2498–2504.

Singh, P. N. P., Shea, C. A., Sonker, S. K., Rolfe, R. A., Ray, A., Kumar, S., Gupta, P., Murphy, P. and Bandyopadhyay, A. (2018). Precise spatial restriction of BMP signaling in developing joints is perturbed upon loss of embryo movement. Development 145.

Somorjai, I. M. L., Martí-Solans, J., Diaz-Gracia, M., Nishida, H., Imai, K. S., Escrivà, H., Cañestro, C. and Albalat, R. (2018). Wnt evolution and function shuffling in liberal and conservative chordate genomes. Genome Biol 19, 98.

Summerhurst, K., Stark, M., Sharpe, J., Davidson, D. and Murphy, P. (2008). 3D representation of Wnt and Frizzled gene expression patterns in the mouse embryo at embryonic day 11.5 (Ts19). Gene Expr Patterns 8, 331–348.

Theiler, K. (1989). The house mouse: Atlas of embryonic development. New York: Springer Verlag.

Tian, A., Benchabane, H. and Ahmed, Y. (2018). Wingless/Wnt Signaling in Intestinal Development, Homeostasis, Regeneration and Tumorigenesis: A Drosophila Perspective. J Dev Biol 6.

Vacik, T., Stubbs, J. L. and Lemke, G. (2011). A novel mechanism for the transcriptional regulation of Wnt signaling in development. Genes Dev 25, 1783–1795.

Vendrell, V., Summerhurst, K., Sharpe, J., Davidson, D. and Murphy, P. (2009). Gene expression analysis of canonical Wnt pathway transcriptional regulators during early morphogenesis of the facial region in the mouse embryo. Gene Expr Patterns 9, 296–305.

Wang, Y., Zhou, C. J. and Liu, Y. (2018). Wnt Signaling in Kidney Development and Disease. Prog Mol Biol Transl Sci 153, 181–207.

Yushkevich, P. A., Piven, J., Hazlett, H. C., Smith, R. G., Ho, S., Gee, J. C. and Gerig, G. (2006). User-guided 3D active contour segmentation of anatomical structures: significantly improved efficiency and reliability. Neuroimage 31, 1116–1128.

Zhao, L., Saitsu, H., Sun, X., Shiota, K. and Ishibashi, M. (2010). Sonic hedgehog is involved in formation of the ventral optic cup by limiting Bmp4 expression to the dorsal domain. Mech Dev 127, 62–72.

